# Evidences of a cytotoxic activity of S-adenosylmethionine on OCI-AML3 cells

**DOI:** 10.1101/004978

**Authors:** K. Aprile von Hohenstaufen, I. Puoti, M. Meloni, B. De Servi

## Abstract

**Background:** The acute myeloid leukemia (AML) cell line OCI-AML3, carrying both NPM1 mutation A and the heterozygous DNMT3A R882C mutation, represents the model for in vitro studies on AML with mutated NPM1^1^. AML with mutated NPM1 harbours a hypo-methylated profile distinct from those of the other AML subtypes^2^. This characteristic is probably related to the inhibitory effect of the mutant DNMT3A on the wild type protein^3^. S-adenosylmethionine (SAM) is a universal methyl donor acting as a coenzyme of DNMT3A. There are growing evidences of the antineoplastic effect of SAM in vitro and in murine models of gastric cancer, colon cancer and hepatocellular carcinoma, where SAM induces the downregulation of several oncogenes^4-10^. Moreover SAM upregulates the expression of DNMT enzymes in lung cancer cells^11^. In our knowledge there are no published data exploring the effect of SAM on the growth of OCI-AML3 cells and its ability to modulate DNMT3A activity in this cell line.

**Study design and methods:** The present data have been generated between August 2013 and April 2014 at the VITROSCREEN facilities in Milan–ITALY. We used a 3-(4,5-dimethylthiazol-2-yl)-2,5-dephenyl tetrazolium bromide (MTT) assay to assess the cytotoxic effect of SAM iodide (Sigma-Aldrich) on OCI-AML3 cells (DSMZ Leibniz Institut). We analyzed then the ability of SAM to induce apoptosis by Tali Image-Based Cytometer (green Annexin V – Alexa Fluor 488 for apoptotic cells, red propidium and green Annexin V-Alexa Fluor 488 for necrotic cells).

**Results:** The MTT assays were performed after having treated the OCI-AML3 cells with various concentrations of the indicated drug for 24 hours. We observed no significant effects on cells viability using 0.5μM, 10 μM and 100 μM of SAM (data not shown). In contrast, a dose dependent cytotoxic effect of SAM on OCI-AML3 cells was evident for concentrations equal or superior to 500 μM, with an IC50 of 500 μM (Figure 1). Since a Cmax of 211(SD 94)μM after single intravenous infusion of SAM was previously reported in healthy voluntarees^12^, we decided to investigate the cytotoxic effect of SAM for concentrations close to 211 μM using the MTT test. A significant dose dependent reduction of the cells viability was observed with SAM 200μM (62,74% viable cells) and SAM 300μM (53.32% viable cells), (Figure 2). The Apoptosis assay after 24 hours of treatment with SAM showed no differences in the percentages of apoptotic cells between the OCI-AML3 cells treated with SAM 300-500-2500 μM and the untreated cells (data not shown). After 72 hours, only a minimal effect on the amount of apoptotic cells was obtained, while a clear dose dependent increase in the proportion of dead cells was noted (Figure 3), confirming the results of the aforementioned MTT tests.

**Conclusions:** SAM showed remarkable in vitro cytotoxic activity on OCI-AML3 cells at concentrations similar to those achievable in humans after intravenous administration. SAM is not able to induce apoptosis of OCI-AML3 cells in vitro after 72 hours of treatment. However, the increase in the amount of dead cells after SAM treatment may be due to mechanisms other than apoptosis. In order to verify if the observed cytotoxicity was mediated by the enzymatic activity of DNMT3A, we planned to repeat the cytotoxicity assays after DNMT3A silencing. The in vivo antineoplastic effect of SAM could be assessed in NOD/SCID mice engrafted with OCI-AML3 cells.

**Authors contribution:** KAvH wrote the study rationale, designed the study, interpreted the data and wrote the article; IP revised the article, MM and BDS planned and interpreted the experiments and BDS performed the experiments.

---

**Figure 1.**
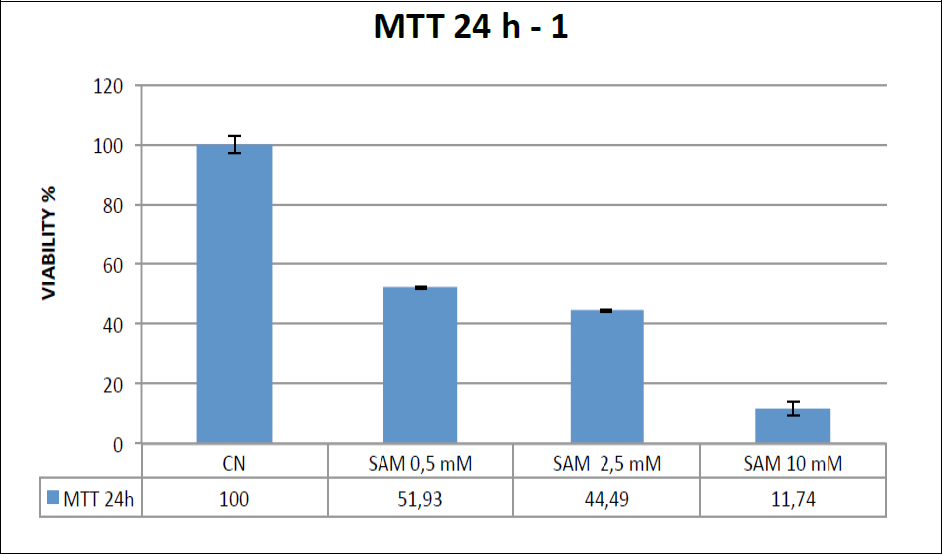
MTT assay after 24 h exposure of OCI-AML3 cells to SAM at 0.5 mM, 2.5 mM and 10 mM. CN= negative control, untreated cells. SAM=S-Adenosylmethionine. Percentages of viable cells are reported. Both SAM treated cells and CN were tested in triplicate samples.

**Figure 2.**
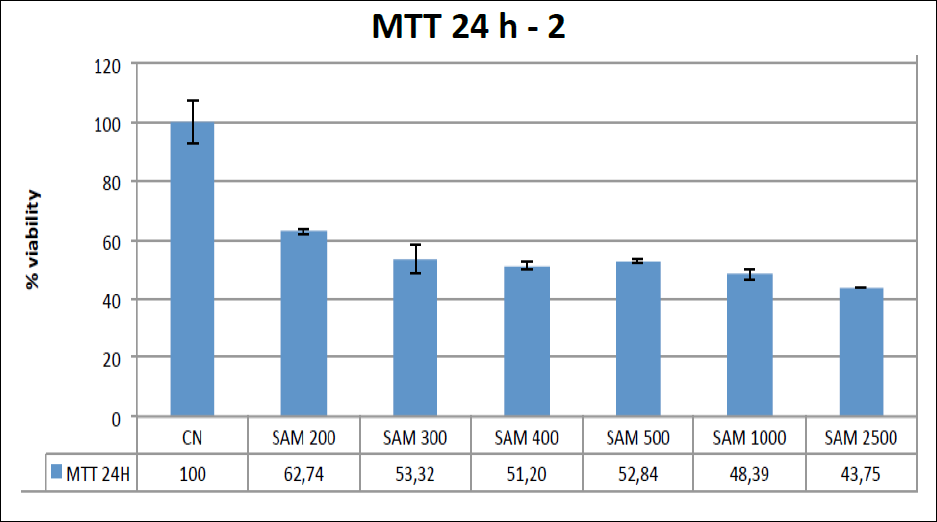
MTT assay after 24 h exposure of OCI-AML3 cells to SAM at 200 μM (0.2mM), 300 μM, 400 μM, 500 μM, 1000 μM, 2500μM. CN=negative control, untreated cells. SAM=S-Adenosylmethionine. Percentages of viable cells are reported. Both SAM treated cells and CN were tested in triplicate samples.

**Figure 3.**
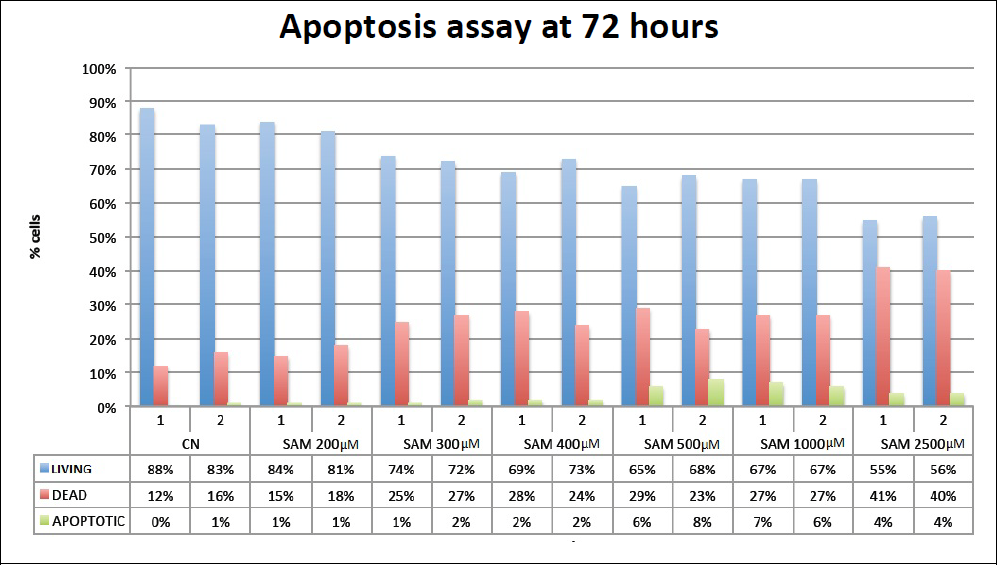
Percentages of Living, Dead and Apoptotic OCI-AML3 cells assessed by Tali image based cytometer after 72 hours treatment with different concentrations of SAM, and after 72 hours in medium (CN=negative control). Both SAM treated cells and CN were tested in duplicate samples.

## References

1. Tiacci E, Spanhol-Rosseto A, Martelli MP, Pasqualucci L, Quentmeier H, Grossmann V, et al. The NPM1 wild-type OCI-AML2 and the NPM1-mutated OCI-AML3 cell lines carry DNMT3A mutations. Leukemia 2012; Mar;26(3):554–7.

2. Cancer Genome Atlas Research Network. Genomic and epigenomic landscapes of adult de novo acute myeloid leukemia. N Engl J Med 2013; May 30;368(22):2059–74.

3. Russler-Germain DA, Spencer DH, Young MA, Lamprecht TL, Miller CA, Fulton R, et al. The R882H DNMT3A Mutation Associated with AML Dominantly Inhibits Wild-Type DNMT3A by Blocking Its Ability to Form Active Tetramers. Cancer Cell 2014; Mar 18;.

4. Chen H, Xia M, Lin M, Yang H, Kuhlenkamp J, Li T, et al. Role of methionine adenosyltransferase 2A and S-adenosylmethionine in mitogen-induced growth of human colon cancer cells. Gastroenterology 2007; Jul;133(1):207–18.

5. Guruswamy S, Swamy MV, Choi CI, Steele VE, Rao CV. S-adenosyl L-methionine inhibits azoxymethane-induced colonic aberrant crypt foci in F344 rats and suppresses human colon cancer Caco-2 cell growth in 3D culture. Int J Cancer 2008; Jan 1;122(1):25–30.

6. Lu SC, Ramani K, Ou X, Lin M, Yu V, Ko K, et al. S-adenosylmethionine in the chemoprevention and treatment of hepatocellular carcinoma in a rat model. Hepatology 2009; Aug;50(2):462-71.

7. Sahin M, Sahin E, Gumuslu S, Erdogan A, Gultekin M. DNA methylation or histone modification status in metastasis and angiogenesis-related genes: a new hypothesis on usage of DNMT inhibitors and S-adenosylmethionine for genome stability. Cancer Metastasis Rev 2010; Dec; 29(4):655–76.

8. Luo J, Li YN, Wang F, Zhang WM, Geng X. S-adenosylmethionine inhibits the growth of cancer cells by reversing the hypomethylation status of c-myc and H-ras in human gastric cancer and colon cancer. Int J Biol Sci 2010; Dec 6; 6(7):784–95.

9. Li TW, Yang H, Peng H, Xia M, Mato JM, Lu SC. Effects of S-adenosylmethionine and methylthioadenosine on inflammation-induced colon cancer in mice. Carcinogenesis 2012; Feb; 33(2):427–35.

10. Li TW, Zhang Q, Oh P, Xia M, Chen H, Bemanian S, et al. S-Adenosylmethionine and methylthioadenosine inhibit cellular FLICE inhibitory protein expression and induce apoptosis in colon cancer cells. Mol Pharmacol 2009; Jul; 76(1):192–200.

11. Ham MS, Lee JK, Kim KC. S-adenosyl methionine specifically protects the anticancer effect of 5-FU via DNMTs expression in human A549 lung cancer cells. Mol Clin Oncol 2013; Mar; 1(2):373–8.

12. Yang J, He Y, Du YX, Tang LL, Wang GJ, Fawcett JP. Pharmacokinetic properties of S-adenosylmethionine after oral and intravenous administration of its tosylate disulfate salt: a multiple-dose, open-label, parallel-group study in healthy Chinese volunteers. Clin Ther 2009; Feb; 31(2):311–20

